# A Visually Interpretable Histopathology-Based Immune Model Predicts T-effector Biology and Response to Immune checkpoint inhibition in Clear Cell Renal Cell Carcinoma Clinical Trial and Contemporary Real-World Datasets

**DOI:** 10.64898/2026.06.21.733614

**Authors:** Averi Perny, Vipul Jarmale, Jay Jasti, Hua Zhong, Alana Christie, Jeffrey Miyata, Aleksandra W. Nielsen, Panayiotis Kontoyiannis, Dinesh Rakheja, Zora Modrusan, Mahrukh Huseni, Ward Kadel, James Brugarolas, Payal Kapur, Satwik Rajaram

**Author notes:** Author of correspondence: Satwik Rajaram, PhD Assistant Professor in Lyda Hill Department of Bioinformatics University of Texas Southwestern Medical Center & Payal Kapur, MD Professor in Department of Pathology University of Texas Southwestern Medical Center. Have contributed equally (Averi Perny, Vipul Jarmale). Conflicts of interest disclosures: None.

## Abstract

Immune checkpoint inhibitors (ICI) are central to the treatment of metastatic clear cell renal cell carcinoma (ccRCC), yet only a subset of patients derive durable benefit, and clinically deployable predictive biomarkers remain an unmet need. RNA-based T-effector signatures capture cytotoxic immune biology and have been associated with ICI response in clinical trial cohorts; however, their clinical implementation is limited by the marked spatial heterogeneity of ccRCC, as well as cost, long turnaround time, sample quality requirements, and limited accessibility. Here, we developed a visually interpretable deep learning (DL) model that predicts a T-cell-enriched immune score directly from hematoxylin and eosin (H&E)-stained whole-slide images. To overcome the inability of H&E morphology alone to distinguish lymphocyte subsets, we trained the model using multimodal spatial supervision from CD8, PAX8, and ERG IHC, which respectively identified cytotoxic T-cell-rich regions, tumor cells, and endothelial cells, thereby constraining immune predictions to relevant tumor microenvironmental niches. The resulting H&E DL Immune score was validated by pathologist review, comparison with held-out CD8 IHC annotations, and independent datasets. The H&E DL Immune score correlated with T-effector RNA scores across independent institutional and IMmotion150 clinical trial cohorts (spearman correlations of 0.726; *p*=5.90x10^-15^ and 0.706; *p*=4.04x10^-19^). As a proof of principle, the score was used to characterize associations with key biological features across large cohorts, including sarcomatoid differentiation, *BAP1* and *PBRM1* mutation status, and additional transcriptomic signatures. In IMmotion150 clinical trial cohort, a median-dichotomized H&E DL Immune score, similar to RNA-based T-effector score, was significantly associated with clinical benefit from atezulumab therapy. In contemporary institutional cohorts of patients treated with frontline ipilimumab plus nivolumab or in initial 3 lines of nivolumab monotherapy, patients in the top quartile of H&E DL Immune score had significantly longer progression-free survival. Collectively, these findings support a scalable and interpretable H&E-based biomarker that captures T-effector biology and can help identify patients with ccRCC more likely to benefit from ICIs.

## Introduction

The management of metastatic clear cell renal cell carcinoma (ccRCC) has been transformed by immune checkpoint inhibitors (ICI), vascular endothelial growth factor receptor tyrosine kinase inhibitors (VEGF-TKIs), and the combination of these agents. Programmed cell death protein 1 (PD-1) inhibitor, Nivolumab (Nivo), improved overall survival (OS) compared with everolimus (a standard of care mammalian target of rapamycin (mTOR) complex 1 inhibitor) in previously treated advanced RCC (1). Dual ICI with Nivo, and the cytotoxic T-lymphocyte antigen 4 (CTLA- 4) inhibitor, ipilimumab (Ipi), has shown an overall response rate (ORR) of ∼40% with 10% complete response (CR) rates (2). Even with combination therapies (either with dual ICI or ICI+ TKI), 40% of the patients fail to respond and neither of these treatments uniformly benefit all patients (2–4). Some patients experience durable responses, whereas others progress early, are exposed to significant toxicity, and financial burden. Thus, there is an unmet need for clinically practical biomarkers that identify the immune state of ccRCC and predict benefit from ICIs.

A range of predictive biomarkers of ICI response have been investigated to date. Two of the most well-studied biomarkers for response to ICI across other tumor types, viz., immunohistochemical (IHC) expression of programmed death-ligand 1 (PD-L1) and tumor mutational burden, have not been demonstrated to have predictive ability in metastatic RCC (2, 5–7). PD-L1, like most IHC based assays, is particularly limited by unclear cutoffs, variability in expression, assay choice, and cell analyzed. However, no correlation has been found between tumor mutational burden and prognostic groups, neoantigen burden, or clinical benefit from ICI (7). *PBRM1* (Polybromo 1) loss has been associated with higher clinical benefit in metastatic ccRCC patients receiving nivolumab (8). However, these patients had received prior VEGF-TKIs, and this association has been questioned by multiple other groups (9). In collaboration with Mayo Clinic, we evaluated the association between BAP1 loss and PD-L1 expression in 1010 ccRCCs and found that unlike tumors with PBRM1 loss, BAP1 deficient tumors are more likely to be PD-L1 positive (17.2% versus 8.3%, p=0.01)(10). Further, data from genetically engineered mouse model (GEMM) showed more lymphocytes (CD4 and CD8 T cell) in *Bap1*-mutated than in *Pbrm1*-mutated mice suggesting a causal relationship. These findings were supported by gene expression profiling data from large tumorgraft (patient-derived xenograft) (11) platform (12) that showed Inflamed RCC subtype to be enriched for *BAP1* mutations (p=7.7x10^-5^) (13).

The most promising work in the field of predictive biomarker come from transcriptomic studies. In IMmotion150, gene expression analyses showed that angiogenesis and CD8 T-effector immune programs were associated with differential benefit from sunitinib versus atezolizumab-containing regimens (7). Subsequent analyses of IMmotion151 and JAVELIN Renal 101 refined the concept that ccRCC is composed of molecular subsets with distinct angiogenic, immune, stromal, metabolic, and proliferative biology, each with potential therapeutic relevance (14, 15). Further these RNA-based biomarkers are now being explored prospectively in the OPTIC RCC trial (NCT05361720) (16). However, RNA-based biomarkers are difficult to incorporate in routine clinical practice due to analytic variability, tumor sampling limitations, turnaround time, high cost, and batch effects. These challenges are particularly relevant in ccRCC, where intratumoral heterogeneity can produce spatially distinct angiogenic and immune microenvironments within the same ccRCC (17–19).

Histomorphology allows for interrogating tumor–environment interactions at a phenotypic level. Indeed, to date, the only practical predictive biomarker is sarcomatoid dedifferentiation in RCC. Multiple studies have shown that sarcomatoid RCC show favorable response to ICI therapy, exhibit PD-L1 expression, and are enriched for *BAP1* loss (7, 20, 21). Interestingly, ccRCC with pancreatic metastases, lack BAP1 loss and are refractory to ICI therapy (11, 22). Broadly, these observations suggest that morphology holds cues for treatment selection for ccRCC patients. Histopathology offers a uniquely scalable biomarker platform because hematoxylin and eosin (H&E)-stained slides are universally available, low cost, and routinely sampled across multiple tumor regions. Recent methods such as HE2RNA (23), DeepPT (24), and SEQUOIA (25) have capitalized on this by training deep learning models to predict transcriptomic programs directly from H&E images using bulk RNA profiles as weakly supervised labels. Despite promising performance, these approaches function largely as black boxes. While prediction heatmaps can localize regions associated with a given transcriptomic signal, they provide limited insight into the specific cellular populations and tissue structures driving those predictions, thereby constraining biological interpretability and clinical trust.

Recent work from our group showed that a visually interpretable H&E-based deep learning (DL) model can predict RNA-based angiogenesis score and anti-angiogenic therapy response in ccRCC, while also providing a pixel-level vascular mask that allows easy, pathologist-facing review (26). This framework addressed a central limitation of conventional weakly supervised models: the inability to understand the cellular basis of the prediction.

Here, we present an interpretable immune model for ccRCC that leverages lymphocyte morphology on H&E to quantify immune infiltration. We first develop and validate the model using IHC-derived ground truth and pathologist review. We then benchmark its performance against IHC and demonstrate its ability to predict T-effector RNA scores across independent institutional and clinical trial cohorts. Next, we apply the model to large archival cohorts lacking matched RNA data, recapitulating established associations between immune infiltration, BAP1 loss, and sarcomatoid differentiation, and illustrating how immune biology can be interrogated directly from H&E at a scale not readily achievable with transcriptomic profiling. Finally, we evaluate the model’s clinical relevance in IMmotion150 and in a contemporary real-world UTSW cohort treated with ipilimumab plus nivolumab and nivolumab monotherapy. Together, these data support the feasibility of deriving a scalable immune biomarker from routine H&E slides and provide a foundation for integrating interpretable H&E-based angiogenic and immune biomarkers in ccRCC.

## Results

### Developing an interpretable multi-cell-type H&E Based Deep Learning Model to quantify immune infiltrate in ccRCC

Because immune-rich tumors are more likely to respond to ICI in ccRCC (7, 14), we set out to quantify immune infiltration directly from H&E (**Figure 1A**). This builds on our H&E DL Angioscore framework (26), in which a model was trained using slide-level bulk RNA and pixel-level IHC. That approach worked for vasculature because vessels are distributed relatively uniformly allowing sampled patches to approximate the whole-slide signal. Immune infiltrate, however, presents two challenges. First, cytotoxic T cells cannot be reliably separated from other small lymphocytes, and lymphocyte nuclei can resemble those of low-grade tumor cells, making immune calls vulnerable to confusion with adjacent tumor cells. Second, immune cells are spatially clustered, often at the tumor-normal interface and in stromal regions rather than distributing evenly across the tumor, Thus, immune content must be assessed across the full tumor region rather than inferred from subsampling. These challenges guided our approach.

**Figure 1.**
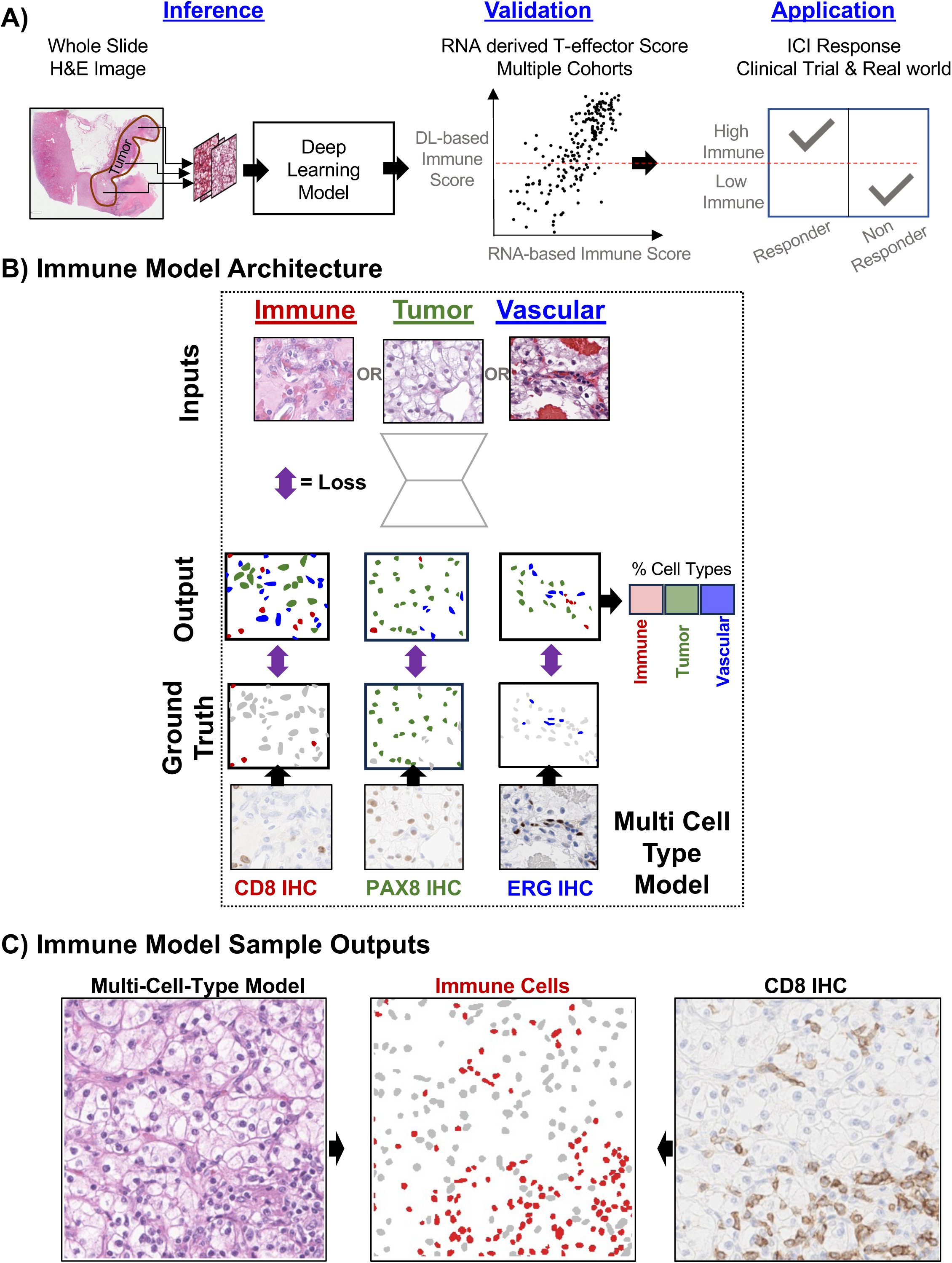
Development and validation of a multi-cell-type deep learning model for immune cell quantification from routine H&E slides. A) Schematic overview of the study workflow. Whole-slide H&E images were analyzed using a deep learning framework to generate an H&E- derived immune score. Model outputs were validated against orthogonal measures of immune infiltration, including immunohistochemistry (IHC) and RNA sequencing–derived T-effector signatures, and subsequently evaluated for association with clinical outcomes and immune checkpoint inhibitor (ICI) response across clinical trial and real-world cohorts. B) Architecture of the multi-cell-type model. The model was trained using aligned H&E and IHC images to simultaneously identify immune, tumor, and vascular cells. Ground-truth labels were derived from CD8, PAX8, and ERG immunohistochemical stains, respectively. By jointly learning multiple cellular compartments, the model generates spatially resolved cell-type predictions and estimates the relative abundance of each cell population within the tissue microenvironment (see methods for details). C) Representative model output. Predicted immune cells from the multi-cell-type model are overlaid on corresponding H&E images and compared with matched CD8 IHC staining. The model accurately identifies regions enriched for lymphocytic infiltration and recapitulates spatial immune patterns observed on immunohistochemistry.

We therefore trained a model to identify immune cells directly on H&E using cell-level ground truth and considered three strategies of increasing complexity. First, we tested whether an off-the-shelf nucleus classifier (HoVer-NeXt)(27), whose lymphocyte class was trained on published annotations from other tissues, could detect immune cells in ccRCC without task-specific training. Second, we trained a custom model using CD8 IHC alone, since the T-effector signature associated with ICI benefit correlates strongly with CD8 T-cell infiltration (7, 14). Using our stain- destain protocol, CD8+ nuclei served as ground truth for a U-Net semantic segmentation model. Because CD8 marks only a subset of immune cells, however, the immune class was defined more broadly than CD8 alone. Third, to reduce confusion between immune cells and morphologically similar tumor and endothelial cells, we added tumor (PAX8) and endothelial (ERG) IHC supervision, yielding a single model that jointly identifies immune, tumor, and vascular cells (**Figure 1B**). Because CD8, PAX8, and ERG were stained on separate sections, each patch carried ground truth for only one marker, so we trained one model across all of them by supervising its output against whichever marker each patch carried (Methods), an approach that to our knowledge is novel. Nuclei themselves were detected with HoVer-NeXt, letting the model focus on classifying cells rather than detecting them.

### Multi-cell model best quantifies immune infiltration

We first compared the three approaches at the single-cell level. On manual review of held-out sections, the multi-cell model’s immune calls visually tracked lymphocytic infiltrate and matched CD8 IHC across a range of immune densities (**Figure 1C, Supplementary Figure 1A**). Quantitatively, single-cell concordance against CD8 was a limited and somewhat misleading benchmark (**Supplementary Table 1**): precision was low for all three models (0.17-0.18) and lowest for the multi-cell model, which at the same time achieved by far the highest recall (0.925). This is expected, since the immune class was intentionally broader than CD8 alone; the model correctly labels other lymphocytes as immune, which appear as false positive when CD8 is used as the only ground truth. Consistent with this, performance improved against CD3, a pan-T marker: precision increased to 0.427 for the multi-cell model, and the multi-cell model led both comparators on all metrics (F1 0.546 vs 0.504 for CD8-only vs 0.392 for HoVer-NeXt; **Supplementary Table 1**). Single-cell ground truth is also intrinsically noisy because of imperfect H&E-IHC registration, cell overlap, segmentation error, and threshold sensitivity, so cell-level concordance understates true performance for all three models.

Because our primary goal was to quantify immune infiltration rather than identify any single cell, we next evaluated performance at the sample level, where aggregation reduces per-cell noise. At the punch level, the multi-cell model’s immune calls correlated strongly with CD3 IHC (Spearman 0.768, p=7.05×10⁻⁹, n=40; **Supplementary Figure 1B**), outperforming both the CD8-only model and HoVer-NeXt (0.50 and 0.156; **Supplementary Table 2**). It also showed the strongest correlation with CD8 IHC in the independent IMmotion150 cohort (Spearman 0.60, p= 1.70×10⁻¹⁰, n=93]; **Supplementary Figure 1C**), again exceeding both comparators (0.51 and 0.28; **Supplementary Table 2**). Because the multi-cell and CD8-only models share the same immune- class definition and nucleus detection and differ only in the added PAX8 and ERG supervision, this improvement reflects the added tumor and vascular context. We therefore adopted the multi-cell model as our immune readout for all subsequent analyses.

### H&E DL Immune score correlates with RNA-derived T-effector biology across independent cohorts

To quantify immune infiltration across a whole slide, we first identified viable tumor on each H&E image using a region classifier, manually reviewed for accuracy (**Supplementary Figure 2A**), and applied the multi-cell model within these regions. In order to minimize the effect of marked spatial heterogeneity of the immune infiltrate (**Supplementary Figure 2B**) we limit our analysis to cases with adequate viable tumor (at least two 1.0 mm cores, approximately 20 mm², per molecular-biopsy adequacy standards; (28, 29)). Additionally, we observed that immune cells frequently concentrated at the tumor-normal interface rather than within the tumor proper. We examined the spatial extent of this peritumoral infiltrate and found it largely confined within 1 mm of the tumor boundary in two independent datasets (**Supplementary Figure 3A).** We therefore defined the <u>H&E DL Immune Score</u> as the density of immune cells across the tumor and a 1 mm peritumoral rim.

To test whether this histologic score reflects underlying immune biology, we compared it to an RNA-derived T-effector score in two cohorts with matched profiling: an institutional UTSW transcriptomic cohort (n=79) and the IMmotion150 clinical trial cohort (n=94). In both, the H&E sections directly flanked the tissue used for RNA extraction, giving tighter spatial correspondence between morphology and expression than is possible in public datasets such as TCGA. The T-effector score was computed from immune effector genes used in prior ccRCC biomarker studies (CD8A, EOMES, PRF1, IFNG, CD274).

The H&E DL Immune score correlated strongly with the RNA T-effector score in both cohorts (Spearman 0.72, p=1.2×10⁻¹³ for UTSW and 0.71, p=1.02×10⁻¹⁵ for IMmotion150; **Figure 2**). It did so more strongly than SEQUOIA (25), a published H&E-to-expression model, in both cohorts (0.716 vs 0.628 for UTSW and 0.711 vs 0.567 for IMmotion150; **Supplementary Table 3**), while also providing a cell-level immune map rather than a region-level expression estimate, so the prediction can be inspected directly on the slide.

**Figure 2.**
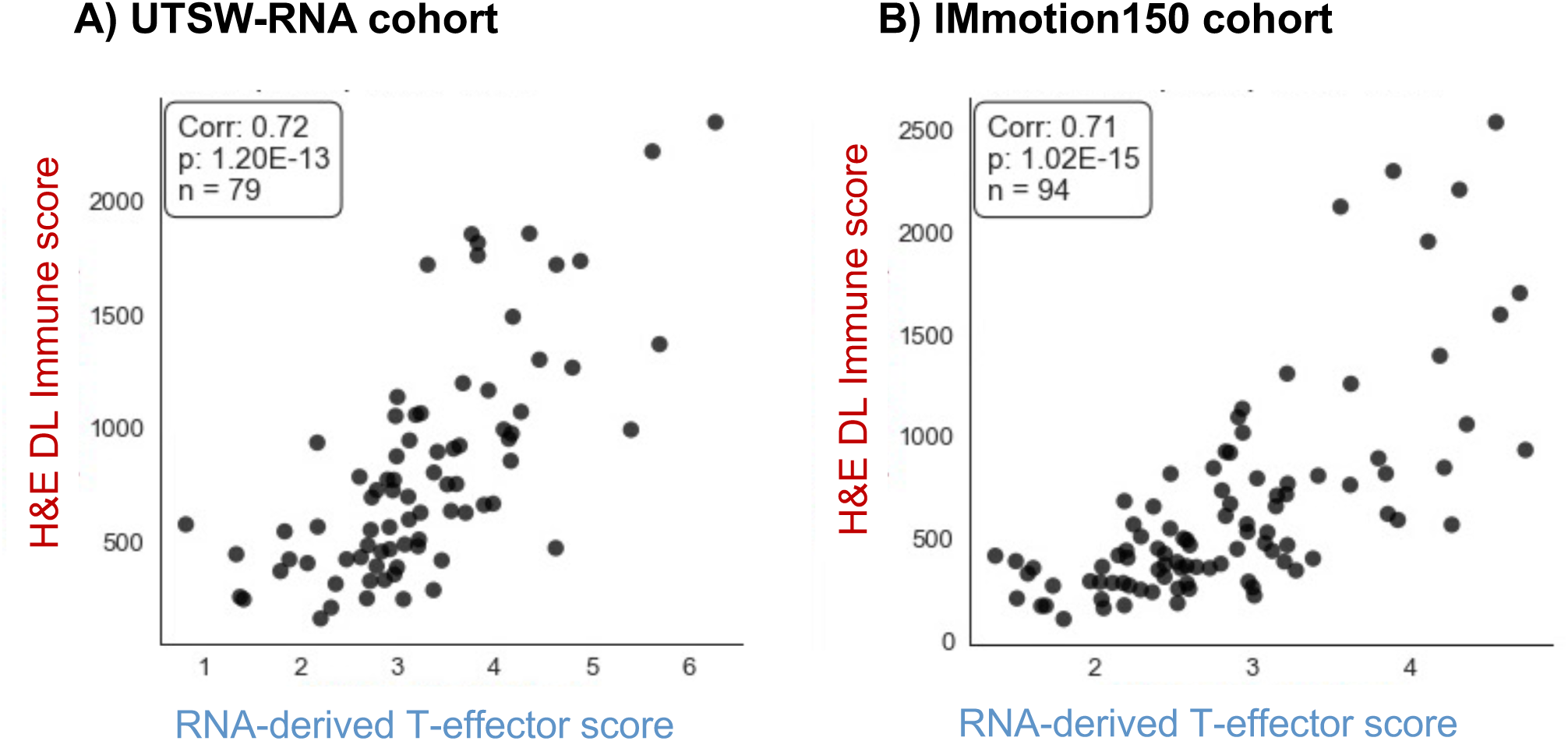
H&E DL Model reliably predicts RNA-based T-effector score across multiple independent cohorts. Each panel shows a scatter plot (each point represents a sample) comparing RNA-based T-effector score (x-axis) and predicted H&E based scores (y-axis). Spearman correlation coefficient along with the p-values and number of samples are displayed in the legend. **A)** Correlation between H&E deep learning immune score and RNA-derived T-effector score in the UTSW-RNA cohort. **B)** Independent validation in the IMmotion150 cohort demonstrating a similarly strong association between model-derived immune infiltration estimates and transcriptomic measures of T-cell effector activity.

This result was not an effected by how the score was defined. Restricting the density to the tumor alone, excluding the peritumoral rim, arguably strengthened correlation with T-effector (**Supplementary Figure 3B**), possibly reflecting the tumor macro-dissection performed for RNA capture. And although the model was trained only on CD8, its score correlated with multiple RNA immune signatures in both cohorts, including the JAVELIN immune signature (15), the ABRS and tertiary lymphoid structure (TLS) signatures (30), and the inflamed ccRCC molecular subset (31), several of which are themselves used to predict immunotherapy response (**Supplementary Figure 3C**). As expected for an immune-specific readout(13), its correlation with the angiogenesis signature and the angiogenic molecular subset was weak or slightly negative (**Supplementary Figure 3C**). The score therefore behaves as a broad immune readout rather than a narrow CD8 count, consistent with the deliberately broad immune class.

Together, these results show that an immune signal read directly from routine H&E, the density of immune cells across tumor and a 1 mm peritumoral rim, approximates transcriptomic T-effector biology in both institutional and trial settings, and does so more accurately and more transparently than a black-box expression predictor.

### Relationship of the H&E DL Immune score with clinicopathologic variables and prognosis

Having validated our H&E based model’s quantification of immune cells, we asked how its predictions correlated with other clinical variables. In the IMmotion151 data, the highest T-effector expression with seen in Cluster 4, a molecular clade that showed the highest PD-L1 expression by IHC (7, 14). We asked if the predictions from the DL model correlated with PD-L1 expression. As shown in **Figure 3A**, H&E DL Immune score highly correlated with PD-L1 expression. Further, cluster 4 was enriched for samples with *BAP1* (BRCA1 associated protein 1) mutations (40%)(14). We and others have reported that *BAP1* and *PBRM1* mutations are largely mutually exclusive (32, 33). Further our observation from *Bap1*-deficient vs. *Pbrm1*-deficeint GEMMs and empirical analyses of tumor microenvironment from PDX models of ccRCC suggests inflamed tumors to be significantly enriched for *Bap1*-deficient tumors unlike *Pbrm1*-deficient tumors that were more angiogenic (7, 13, 34–37). To explore this phenotype, we applied the Immune model to our large, published cohort with WSI and available BAP1 and PBRM1 IHC status (38). Consistent with prior findings, we found H&E DL Immune score to be significantly higher in ccRCC with BAP1 loss as compared to those with PBRM1 loss (**Figure 3B**; *p*=5.202 x10^-04^). These findings were validated in the TCGA cohort with *BAP1* and *PBRM1* gene status (**Figure 3B**; *p*=5.49 x10^-02^).

**Figure 3.**
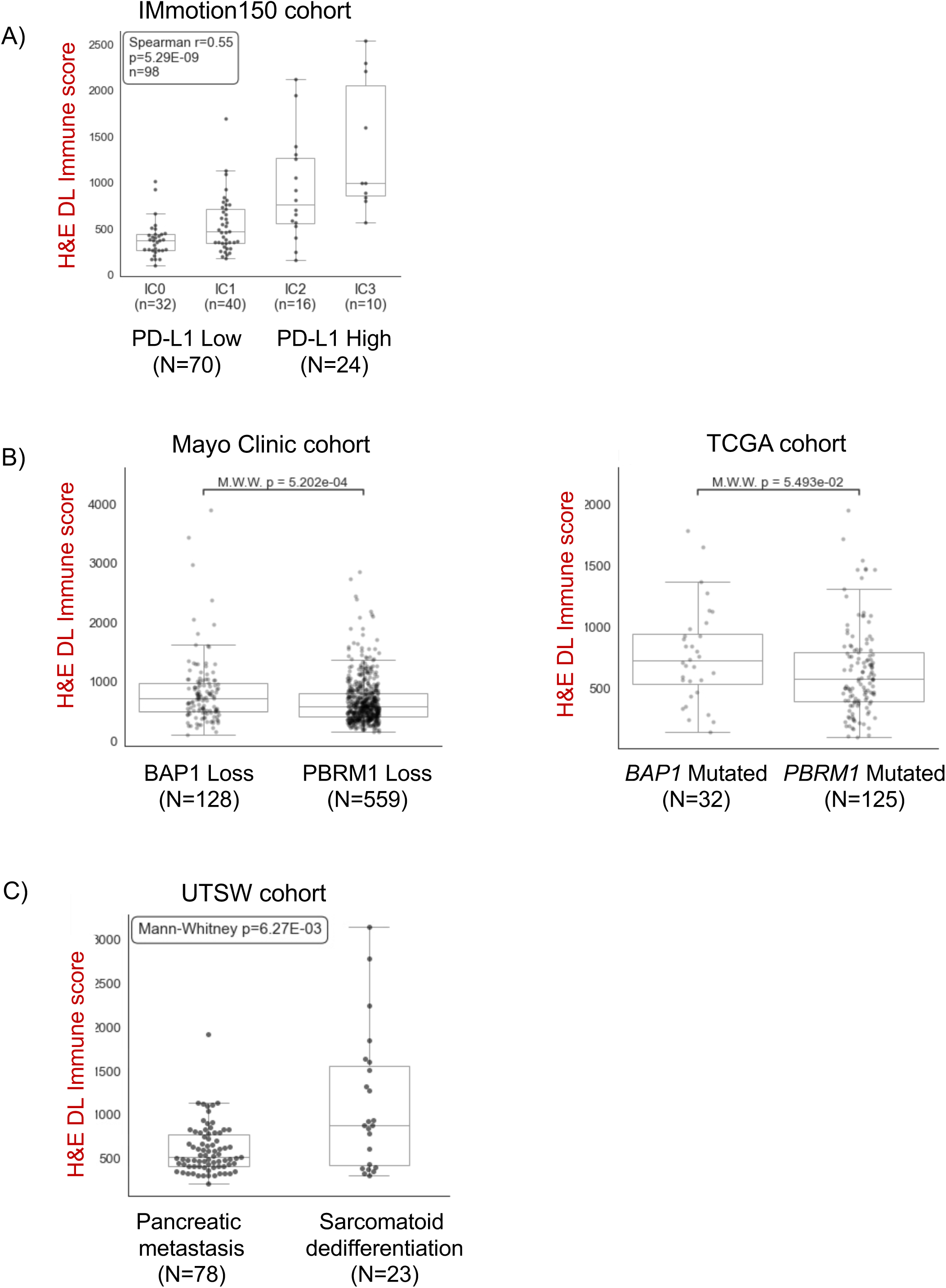
Model predictions correlate with biologically relevant molecular and clinical features across independent cohorts. The H&E DL Immune score model was applied to multiple independent datasets and its output was compared to various clinical variables. **A)** Correlation of H&E immune score with reported PD-L1 IHC expression (IC0-IC3) in the IMmotion150 cohort. As expected, increasing immune score was seen with higher PD-L1 expression. Correlation was assessed using Spearman rank correlation. **B)** H&E immune scores according to BAP1 and PBRM1 status in Mayo Clinic (by IHC) and the TCGA (by mutation analyses) cohorts. Tumors harboring BAP1 alterations exhibit higher immune infiltration, consistent with prior reports linking BAP1 loss to inflammatory tumor microenvironments (rare cases with loss of both BAP1 and PBRM1 are considered as exhibiting BAP1 loss). Statistical significance was assessed using the Mann–Whitney–Wilcoxon test. **C)** Association of H&E immune score with sarcomatoid dedifferentiation in the UTSW cohort. As reported previously, sarcomatoid dedifferentiation demonstrates substantially higher immune infiltration. This contrasts with tumors that metastasize to pancreas.

The significance of these molecular subtypes is also illustrated by our recent study on ccRCC patients with pancreatic metastases (11, 39), where we found that the ccRCC that metastasize to the pancreas lack BAP1 mutations. This contrasts with ccRCCs with a sarcomatoid component that traditionally have poor prognosis (40) but are highly sensitivity to ICI and enriched for BAP1 loss (41, 42). Given these findings we leveraged available UTSW WSI datasets and identified ccRCC with sarcomatoid dedifferentiation (n=23) and compared them to those from patients with pancreatic metastasis (n=78). As shown in **Figure 3C** ccRCC with sarcomatoid dedifferentiation showed significantly higher H&E DL Immune score compared to pancreas metastasis (*p*=6.27 x10^-03^).

We leveraged the publicly available TCGA cohort and did not find significant correlation between the H&E DL Immune score and the World Health Organization/ International Society of Urological Pathology (WHO/ISUP) nucleolar grade, TNM stage, or tumor size (**Supplementary Figure 4A- C**). Consistently, Kaplan-Meier curves stratifying patients based on their H&E DL Immune score using quartile cutoffs did not reveal significant correlation between Immune score and overall survival (**Supplementary Figure 4D**). Next, we tested the extent to which the H&E DL Immune score differed between samples from primary tumor and metastatic sites. We did not observe significant difference in H&E DL Immune score based on sample site (**Supplementary Figure 4E-F)**.

Overall, our results show that that DL techniques are powerful techniques and biomarkers such as H&E DL Immune score can be successfully leveraged to explore biology; and illustrate that ccRCC with high Immune scores are associated with high PDL1 IHC expression, sarcomatoid dedifferentiation and BAP1 loss.

### H&E DL Immune score predicts ICI therapy response in IMmotiton150

Given that RNA-derive T-effector program is a predictor of response to ICI therapy, we next sought to test the predictive performance of our H&E DL Immune score. First, we applied our model to IMmotion150 clinical trial. This phase 3 trial evaluated atezolizumab alone, atezolizumab plus bevacizumab, and sunitinib in treatment-naïve metastatic RCC and has served as a foundational dataset for angiogenesis and T-effector biomarker discovery in ccRCC. Although atezolizumab-based regimens are not contemporary standard-of-care regimens for ccRCC, IMmotion150 provides a rigorously annotated clinical trial cohort with matched H&E slides, RNA expression data, and clinical outcomes. Using this dataset, we compared the predictive value of the H&E DL Immune score to that of the RNA-derived T-effector score and CD8 IHC for response to atezolizumab. Following the original IMmotion150 publication we stratified the patients into low/high Immune groups based on the median value of each assay we examined the relationship to Progression free survival (PFS), and generated Kaplan Meir curves and performed Cox-proportional hazards calculations (DL model: HR 0.35, *p*=0.005; T-effector: HR 0.32, *p*=0.003; **Figure 4A).** The H&E based predictions were comparable to RNA-based predictions of atezolizumab response and slightly superior to the CD8 IHC. This point was further reinforced by separate analyses comparing our three assays in terms of a) the fraction of patients who responded to atezolizumab among high/low Immune groups (**Supplementary Figure 5A-B**) and b) the AUC in predicting the objective response (responder or not) based on Immune cell density (**Figure 4B**).

**Figure 4.**
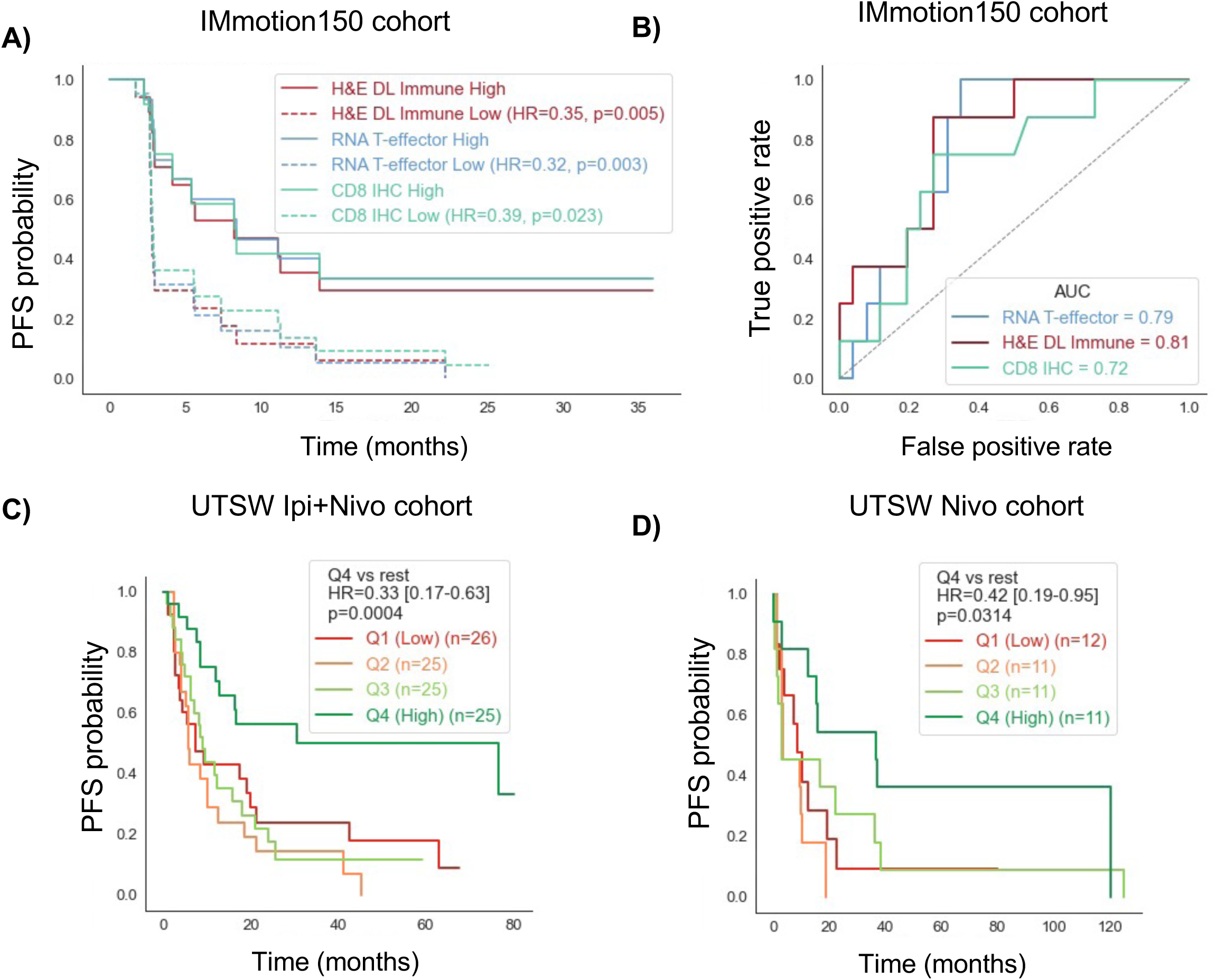
H&E DL Immune score predicts clinical benefit from immune checkpoint blockade. **A)** Progression-free survival stratified by H&E immune score in the IMmotion150 immunotherapy-treated cohort. Comparisons with RNA-derived T-effector score and CD8 IHC are shown. Patients were stratified at the full-cohort median of each marker; A and B use the 34 atezolizumab-treated patients with H&E immune score, RNA T-effector, and CD8 IHC all available. **B)** Receiver operating characteristic analysis demonstrating predictive performance of the H&E immune score for treatment benefit relative to molecular biomarkers. **C)** Progression- free survival in the real-world UTSW cohort treated with frontline ipilimumab–nivolumab stratified by H&E immune score. Patients are stratified based on H&E DL quartile score and the highest quartile are compared with others. **D)** Similar Progression-free survival in patients treated with nivolumab monotherapy in any line. Survival curves are Kaplan-Meier estimates; hazard ratios are from univariate Cox models (high vs low group) and p-values are from two- sided log-rank tests.

### H&E DL Immune score predicts benefit from contemporary ICI regimens in real-world cohorts

To determine whether the H&E DL Immune score has predictive value in contemporary clinical practice, we evaluated a UTSW cohort of patients with metastatic ccRCC treated with ICI regimens currently relevant to clinical care. The primary contemporary cohort included patients treated with frontline ipilimumab plus nivolumab (n=101) between 2019 to 2024. A second cohort included patients treated with nivolumab monotherapy (n=45) in first three lines of therapy. As multiple WSI were available per patient in these cohorts, we summarized each patient by the maximum H&E DL Immune score across their slides, analogous to histologic grade, which is assigned from the highest-grade focus rather than an average (see Methods).

In the frontline ipilimumab plus nivolumab cohort, high H&E DL Immune score was associated with improved PFS with a c-index of 0.592 (**Supplementary Table 4**). Using the median value to stratify, patients with high scores had longer PFS than those with low scores (**Supplementary Figure 6A**) obtained a HR of 0.55 (95% CI: 0.35-0.88) with a p value of 0.012. Interestingly, best stratification with hazard ratio of 0.33 (95% CI: 0.17-0.63) with a p value of 0.0004 was obtained using the highest quartile threshold H&E DL Immune score (**Figure 4C**).

In the nivolumab monotherapy cohort, high H&E DL Immune score was also associated with improved outcome (**Supplementary Figure 6A**, **Supplementary Table 4**. HR 0.51, 95% CI 0.26– 1.02, p = 0.051, c-index 0.556). In the quartile-based analysis, the top immune-score quartile again showed improved outcome compared with the rest of the cohort (**Figure 4D**; quartile 4 vs rest HR 0.42, 95% CI 0.19–0.95, p = 0.031).

We asked if the results would be influenced if we randomly selected a WSI. As shown in **Supplementary Figure 6A,** bootstrap analysis by selection of a random slide did not significantly affect the correlation with PFS when a pooled or most inflamed slide was selected. These results indicate that an H&E-derived immune score can identify ccRCC tumors more likely to benefit from PD-1-based ICI therapy in a contemporary real-world setting. The observation that the model performs in both ipilimumab plus nivolumab and nivolumab monotherapy cohorts supports the hypothesis that the H&E DL Immune score captures a shared T-cell-inflamed biology relevant to immune checkpoint blockade.

## Discussion

In this study, we developed and validated a visually interpretable H&E-based immune model for ccRCC. The model uses routine H&E whole-slide images to generate an immune score that correlates with CD3 IHC, approximates RNA-derived T-effector biology, and associates with benefit from ICI therapy in both a clinical trial dataset and a contemporary real-world cohort. These findings extend our prior H&E DL Angioscore framework from angiogenic biology to immune biology and support the feasibility of using routine pathology slides to derive clinically relevant, spatially informed biomarkers in ccRCC.

A key feature of this work is the multi-cell-type model architecture. Immune quantification from H&E is inherently more challenging than vascular quantification because CD8-positive T cells cannot be reliably separated from CD4-positive T cells or other small lymphocytes by morphology alone. A model trained only on CD8 IHC may therefore learn a nonspecific lymphocyte signal, other cell types or may be confounded by tumor nuclei of low-grade tumors. By incorporating PAX8-defined tumor epithelium and ERG-defined vascular compartments, the model is encouraged to interpret immune infiltrates within the relevant tissue context. The improved performance of the multi-cell-type model compared with the CD8-only model supports this design principle.

The model’s correlation with RNA T-effector scores is particularly significant. Prior clinical trial analyses have shown that T-effector and angiogenesis programs provide complementary information about therapy response in metastatic ccRCC (7, 14, 15). However, transcriptomic signatures have not been widely adopted clinically because of cost, turnaround time, batch effects, tissue requirements, and limited spatial representation. An H&E-based immune model could overcome several of these barriers. H&E slides are ubiquitous, can be analyzed rapidly, and often represent multiple tumor regions.

The output of our model was strongly predictive of the RNA-derived T-effector score across multiple cohorts, with Spearman correlation approaching 0.75 on multiple real world and clinical trial datasets (UTSW/IMmotion150). Our H&E DL Immune score allowed us to explore the relationship of T-effector biology with various variables on a large cohort without the need for RNA-seq analysis. Our analysis reinforces the notion that in ccRCC immune infiltrate inversely correlated with tumor angiogenesis, is enriched in tumors with sarcomatoid change and tumors with BAP1 loss compared to those with PBRM1 loss. Finally, in the IMmotion 150 clinical trial data, we found that the ability of our model to predict response to ICI therapy rivaled that of the RNA.

The distinguishing aspect of our DL model is that it can provide visually interpretable predictions and thereby overcome its limitation as a “black box”. This is critical for quality evaluation and clinical adoption. In multiple cancers, previous efforts have demonstrated prediction of specific genes and pathways from H&E images (23, 43–46).

Our findings also highlight the need to distinguish biological validation from clinical validation. IMmotion150 provides an invaluable dataset because it contains pretreatment H&E slides, RNA expression data, and clinical outcomes across treatment arms. However, atezolizumab-based regimens are not current standard therapy for ccRCC. For this reason, we used IMmotion150 primarily as a trial-based biological validation cohort and then tested clinical relevance in UTSW patients treated with contemporary PD-1-based regimens, including frontline ipilimumab plus nivolumab and nivolumab monotherapy. The association of high H&E DL Immune score with improved outcomes in these real-world cohorts strengthens the argument that the model captures clinically meaningful immune biology.

This work has several potential clinical implications. First, an H&E-based immune score could help identify patients more likely to benefit from ICI-based therapy, particularly when RNA testing is unavailable or impractical. Second, because the model is spatially resolved, it could quantify intratumoral and peritumoral immune heterogeneity across multiple slides, biopsies, or tumor regions. Third, the immune model could be integrated with the H&E DL Angioscore to build a composite histopathology-based biomarker that distinguishes angiogenic, immune-inflamed, mixed, and immune-cold tumors. Such a biomarker may eventually help guide selection among ICI/ICI, ICI/VEGF-TKI, VEGF-TKI-dominant, or investigational approaches.

The study also has limitations. The model was trained using CD8 IHC, but H&E morphology cannot specify CD8 lineage with certainty; therefore, the H&E DL Immune score should be interpreted as a T-cell-enriched immune score rather than an exact CD8 count. Validation against CD3 IHC and RNA T-effector signatures helps address this limitation but does not fully resolve the underlying cellular composition. The UTSW treatment cohorts are retrospective and may include heterogeneity in specimen timing, prior therapy, tissue site, and line of therapy. IMmotion150 provides trial validation but uses regimens that are not contemporary standards of care. Thresholds used for dichotomization require prospective locking and external validation. Finally, immune infiltration is spatially heterogeneous, and biopsy samples may undersample immune-rich or immune-poor regions despite minimum tissue adequacy criteria.

Future work should focus on external validation in independent cohorts treated with ipilimumab plus nivolumab, nivolumab monotherapy, and ICI/VEGF-TKI combinations. Ideally, the model should be evaluated in clinical trial datasets with pretreatment H&E slides, response annotation, and RNA or spatial immune profiling. Additional studies should assess whether the H&E DL Immune score adds predictive value beyond IMDC risk group, sarcomatoid differentiation, PD-L1 expression, CD8 IHC, and RNA T-effector signatures. Finally, integration with H&E-based angiogenesis, necrosis, tumor architecture, and radiologic features may yield a more comprehensive model of therapy response in ccRCC.

In summary, we present an interpretable H&E-based immune model that captures T-effector biology and predicts benefit from immune checkpoint blockade in ccRCC. By converting routine pathology slides into spatially resolved immune biomarkers, this approach may bridge molecular biomarker discovery and practical clinical implementation.

## Methods

### Study Cohorts

All cohorts consisted of formalin fixed paraffin embedded (FFPE) hematoxylin and eosin (H&E) stained whole slide images (WSIs) scanned at either 20X (∼0.5 microns per pixel) or 40X (∼0.25 microns per pixel) magnification. Because the deep learning model was developed using 20x images, all 40x images were downsampled by a factor of two prior to analysis. For cohorts used to evaluate the final multicell deep learning (DL) immune score, slides were excluded if they lacked sufficient tumor tissue for analysis (<20 mm^2^ tumor area) or represented lymph node metastases, which precluded reliable assessment of peritumoral regions. Detailed cohort assembly and exclusions are provided below.

1. UTSW CD8 training dataset: The primary training dataset for CD8+ T-cell segmentation consisted of 19 ccRCC WSI that capture a broad range of ccRCC tumor grade and morphologic patterns. These slides were initially stained with H&E, imaged at 20X magnification (Aperio scanners), subsequently de-stained, re-stained with antibody for CD8 (clone C8/144B; Agilent CA) and re-imaged. Fifteen WSI were used for training and 4 were held out for testing.
2. UTSW ERG training dataset: The endothelial-cell training dataset consisted of 7 ERG- restained ccRCC WSIs generated using the same stain-destain workflow described above and stained with ERG (clone 9FY; Biocare CA) immunohistochemistry (IHC). All 7 WSIs were used to identify endothelial-cell nuclei during multicell model development.
3. UTSW PAX8 training dataset: Tumor-cell ground truth was generated from six PAX8- restained tissue microarray (TMA) sections (593 punches) generated using the same stain- destain workflow described above and stained with PAX8 (clone MRQ-50; Cell Marque CA) IHC. Punches with tissue folds/loss/damage, poor PAX8 staining, or failed registration were excluded, leaving 139 evaluable punches. All 139 punches were used to identify tumor-cell nuclei during multicell model development.
4. UTSW CD3 validation dataset: To validate immune cell identification against a pan-T-cell marker, CD3 IHC was performed on two TMA sections (96 punches) from a previously described ccRCC cohort (33, 38). Slides were stained with H&E and imaged, then destained, restained with antibody for CD3 (clone F7.2.38; Agilent CA), and re-imaged. Following quality check, 56 punches were excluded due to tissue that was missing, folded, or otherwise non-evaluable, leaving 40 adequately registered tumor-containing punches.
5. UTSW Sequenced (Seq) cohort: This cohort was used to assess the relationship between H&E DL Immune Score and RNA T-effector score. This cohort comprised 79 ccRCC H&E WSI from 79 patients with matched clinical RNA sequencing (Caris Life Sciences). RNA-based T-effector scores were computed as described below.
6. TCGA KIRC cohort: This cohort served as our primary external dataset for clinicopathologic analyses (tumor grade, stage, and overall survival) and was also used for relating the immune score to BAP1/PBRM1 mutation status. H&E WSIs were downloaded from the NIH Genomic Data Commons (47–49); grade, stage, survival, and BAP1/PBRM1 mutation calls were obtained from the cBioPortal PanCancer Atlas (48). We restricted to ccRCC cases with a single diagnostic slide per patient, excluded low-quality, non-ccRCC, and duplicate slides identified using the criteria from our prior work (26), and applied the tumor-area cutoff, leaving 439 evaluable WSIs (most 40X downsampled to 20X, a minority native 20X). Each analysis used the subset with the relevant annotation, with group sizes reported in each figure.
7. Mayo Clinic cohort: This cohort was used to evaluate the impact of loss of BAP1 and/or PBRM1 on the immune infiltrate. We used a subset of a previously published (38) ccRCC cohort from the Mayo clinic of H&E-stained WSIs (scanned at 20X) with matched BAP1 and PBRM1 status, assessed by IHC as wildtype, Universal Loss, or Localized Loss (loss confined to a region of the slide). Among slides with adequate tissue, 106 showed universal BAP1 loss, 559 showed universal PBRM1 loss, and 22 showed simultaneous universal loss of both, which were grouped with BAP1 loss for analysis; an additional 7, 22, and 0 slides in these respective classes were excluded for inadequate tissue.
8. IMmotion150 cohort: The IMmotion150 clinical trial cohort was used to evaluate associations between the H&E-derived DL immune score, and reported RNA-based T-effector score, CD8 IHC, and response to atezolizumab. Following exclusion of poor quality (n=5), duplicate WSI (n=3), lymph node metastases (n=2), and WSIs failing tissue adequacy criteria (n=32), 98 evaluable WSIs remained. Among these, RNA expression data were available for 94 cases and CD8 IHC data for 93 cases. Response analyses were restricted to patients treated with atezolizumab (n=36) and modality comparisons were performed using cases available across all relevant modalities. Of note, this dataset is a subset of 305 patient data reported in the IMmotion150 study (7).
9. UTSW Ipilimumab plus Nivolumab (Ipi+Nivo) cohort: This is our principal real-world cohort for assessing the relationship between our H&E DL Immune Score and response to immune checkpoint inhibitors. It comprises 101 patients with metastatic ccRCC treated at UTSW between 2018 and 2024. Unlike the IMmotion150 trial cohort, in the UTSW response cohorts (Ipi+Nivo and Nivo), each patient contributed multiple H&E slides. Thus the 101 patients were represented by 230 WSI meeting the tissue adequacy criteria. Of these, 192 slides are from primary tumors and 38 from non-lymph-node metastases, with patients contributing 1 to 4 slides each (7, 68, 17, and 9 patients with 1, 2, 3, and 4 slides, respectively). Patients were identified through the Kidney Cancer Explorer (KCE), an IRB-approved institutional clinicopathologic database (50). Slides were selected to capture the largest and most representative areas of highest-grade tumor present in each case. Progression-free survival (PFS) was abstracted from the electronic medical record by investigator (PK) blinded to model outputs.
10. UTSW Nivolumab (Nivo) cohort: As an additional real-world ICI cohort, this dataset comprises 45 patients with metastatic ccRCC who received nivolumab monotherapy within the first three lines of systemic therapy at UTSW, represented by 74 WSI meeting adequacy criteria. Of these, 66 slides are from primary tumors and 8 from non-lymph-node metastases, with patients contributing 1 to 3 slides each (25, 11, and 9 patients with 1, 2, and 3 slides, respectively). Similar to Ipi+Nivo cohort, patients were identified through the KCE using the same approach described above, and up to three representative tumor-containing slides per patient were retained, prioritized by highest tumor nuclear size. PFS was abstracted from the electronic medical record blinded to model outputs.
11. UTSW sarcomatoid and pancreatic metastasis cohort: To explore biologically distinct disease states, separate cohorts of sarcomatoid ccRCC and pancreatic metastases were assembled from KCE-linked pathology archives. The sarcomatoid cohort comprised 23 WSIs (20 primary and 3 metastatic) from 21 patients with pathologist-confirmed sarcomatoid differentiation; consistent with the region-identification approach described above, the immune score was computed only over tumor regions with sarcomatoid morphology, which were manually defined when other morphologies were present on the slide. The pancreatic metastasis cohort comprised 78 WSIs from 18 patients with previously reported pancreatic metastatic disease, comprising 55 primary tumor and 23 metastasis WSIs (pancreatic or other sites) (11).

### RNA Immune score calculation

For the UTSW-Seq cohort, the raw transcriptome sequencing data was processed by the SCHOOL (51) with human reference genome version GRCh38.86 and Fragments Per Kilobase of transcript per Million mapped reads (FPKM) genes were generated. FPKM was normalized to Transcripts Per Kilobase Million (TPM), then log-transformed with 1 added to avoid taking log of zero. For other cohorts, namely TCGA and IMmotion150, TPM data was directly obtained. The T-effector signature genes for determining Immune score are *CD8A*, *EOMES*, *PRF1*, *IFNG*, and *CD274* following the IMmotion150 and IMmotion151 studies (7, 14). The Immune score for each tumor sample was computed by the mean log transformed TPM of the Immune signature genes.

Additional transcriptomic signatures were derived from previously published gene sets, including the atezolizumab–bevacizumab response signature (ABRS)(52) from Zhu et al., ccRCC molecular subtype signatures (ccrcc1–4)(31) from Beuselinck et al., tertiary lymphoid structure (TLS) signatures (30) from Amisaki et al., the tumoral low HLApr (tLHP) signature (53) from Kinget et al., and the JAVELIN immune and angiogenesis gene signatures from Motzer et al.(15). An interleukin (IL) signature was defined using IL6, IL7, and IL8. For all signatures, expression values were transformed as log2(TPM + 1), normalized to gene-wise z-scores, and the signature score for each sample was calculated as the mean z-score across all genes in the corresponding signature. Among the ccrcc4 signature genes, expression of IGHA1, IGHD, IGKC, and IGKV1-5 were not available in the IMmotion150 cohort and were therefore excluded from the score calculation.

### Training data generation

The H&E DL Immune model was trained on H&E image data paired with nucleus-level, single-marker cell-type labels. Three separate re-stain datasets provided ground truth for the three compartments: CD8 for immune (T) cells, ERG for endothelial/vascular cells, and PAX8 for tumor cells. In each, the marker IHC was registered to the matched H&E, binarized to a positive/negative stain mask, and intersected with a nuclear segmentation of the H&E so that every segmented nucleus received a positive or negative call for that marker. These per-nucleus, single-marker labels served as ground truth for the corresponding class of the multi-cell-type H&E model (see Model). The same overall sequence was used throughout; the nuclear segmenter and the overlap rule differed by marker, as below.

CD8 and ERG (whole-slide images):

1. Re-staining and region selection: as described above adequately registered regions of tumor tissue, free of artifacts were annotated for patch extraction.
2. H&E–IHC registration: each H&E and IHC pair was aligned in two steps, a manual rigid slide-level alignment in QuPath (54) followed by automatic non-rigid registration of the shared hematoxylin channels using SimpleElastix’s multi-resolution pyramid framework (v2.0.0rc2 (55), affine then B-spline deformable as described previously (26). Registration used 512x512px patches, from which a centered 416x416px pair was extracted at 20X to remove edge effects.
3. IHC binarization (pixel level): a U-Net classifier was trained to separate marker-positive DAB staining from negative staining and non-specific/artifactual DAB, then applied to the IHC patches to produce a binary positive/negative pixel mask (artifact pixels treated as negative).
4. Nucleus-level assignment: nuclei were segmented from the H&E using HoVer-NeXt (lizard_convnextv2_large)(27); a nucleus was labeled marker-positive if more than 50% of its area overlapped the positive pixel mask, and negative otherwise.

#### PAX8 (TMA)

PAX8 ground truth followed the same logic on TMA sections, with three changes forced by the material and staining. Each section was split into individual punches using QuPath’s TMA layout detection, and each H&E/IHC punch pair was registered with SimpleElastix as above; punches with tissue damage, poor PAX8 staining, or failed registration were excluded (see Cohorts). First, because PAX8 staining quality varied across punches and the U-Net did not generalize, the IHC was binarized with a QuPath pixel classifier [v0.2.3] rather than the U-Net. Second, nuclei were segmented with Hover-Net (PanNuke, tiatoolbox v1.4.0). Third, because PAX8 staining is grainier and the pixel classifier tends to under-predict, a nucleus was labeled PAX8-positive when at least 15 pixels overlapped the positive mask (at 20X), rather than the 50% rule used for CD8/ERG.

#### Data splits

the CD8 dataset included 4 slides (out of 19) held out for evaluation; the ERG and PAX8 datasets were used in full for model development.

#### CD3 Validation

To validate immune-cell identification against pan-T-cell ground truth, the H&E DL Immune model’s immune calls were compared to CD3 IHC on a TMA (cohort and punch counts in Cohorts). CD3 ground truth was constructed as for the training markers (Training data generation): the CD3 IHC was registered to the matched H&E, binarized to a CD3-positive mask, and intersected with the HoVer-NeXt nuclear segmentation so that each nucleus received a CD3 label (based on a majority vote). Two steps differed from the training pipeline: registration used palom (56) (a coarse affine alignment followed by block-wise affine refinement) rather than the QuPath/SimpleElastix pipeline, and the CD3-positive mask was taken from a QuPath positive pixel classifier of the IHC. The immune calls were assessed against this ground truth at two levels. At the single-cell level, each nucleus’s immune call was compared to its CD3 label across all punches, and precision, recall, and F1 were computed, with nuclei the model did not call excluded so the metric reflects classification of detected nuclei. At the punch level, the proportion of nuclei called immune was correlated with the proportion called CD3-positive (Spearman).

### Model

The multi-cell-type H&E model was a U-Net with a ResNet-18 encoder that performed pixel-wise semantic segmentation of each 416x416px H&E patch (20X) into five classes: background, tumor, immune, endothelial, and other. The “other” class absorbed cells outside the three labeled compartments, including non-T-cell immune cells. The network produced a per-pixel softmax over the five classes, which were resolved to individual nuclei at inference (see Quantification).

### Model Training

Training used the H&E patches with per-nucleus ground-truth masks from the CD8, ERG, and PAX8 datasets described above. The central constraint is that ground truth was available for only one marker per patch: each patch carried a three-class mask (background, marker-positive, marker-negative) for its own stain, and no patch was labeled for all compartments at once. To train a single five-class model under this constraint, predictions were mapped into each patch’s marker-specific three-class space before the loss was computed. For a given marker, the background output mapped to background, the output class corresponding to that marker mapped to marker-positive, and the remaining output classes mapped to marker-negative; for example, on a PAX8 patch the tumor output is the positive class while immune, endothelial, and other are all negative. The single exception was the immune output on CD8 patches, which was assigned equally to marker-positive and marker-negative, reflecting that CD8 marks only the T-cell subset of immune cells. The loss was evaluated in this three-class space, so the model was supervised only on distinctions the corresponding stain could verify, and the five output classes were learned jointly across the three datasets rather than from any fully labeled image.

The objective was a weighted combination of a custom negative log-likelihood term (implemented directly, as the standard cross-entropy was unstable in backpropagation here) and a Dice loss, combined as 0.2 and 0.8 respectively and computed on the mapped predictions. Two weighting schemes addressed imbalance: per-marker class weights, because the marker-positive class was rare relative to negative and background, and per-marker sample weights, so the larger CD8 dataset did not dominate the smaller ERG and PAX8 sets. Patches were augmented with horizontal and vertical flips and light HED color jitter. The model was trained for 75 epochs using Adam (learning rate 1e-4, batch size 5), saving checkpoints every five epochs.

### Nucleus Level Inference

Our trained model is fundamentally a semantic segmentation (pixel classification) rather than a nucleus classification model. To produce nucleus-level calls, we used HoVer-NeXt (lizard_convnextv2_large) (27) to identify individual nuclei, and each nucleus was assigned to the class with the most pixels assigned to it by our model. Nuclei whose majority class was background (not called a nucleus by the model) were dropped from downstream analysis.

### Comparator Models

**CD8-only model**: the multi-cell-type model with the tumor and endothelial output classes removed, trained on the CD8 re-stain data alone (three output classes: background, immune, other). The immune compartment is defined identically to the multi-cell-type model; architecture and training procedure otherwise followed the full. Nucleus level inference was performed just as with our full model, except with a reduced set of classes. Since the underlying nucleus identification is the same, comparing this model against the full model isolates the contribution of the auxiliary PAX8 and ERG supervision to immune-cell identification.

**HoVer-NeXt**: a public off-the-shelf nuclei segmentation and classification pipeline (27). We applied the authors’ lizard_convnextv2_large model using the provided inference code. We took its lymphocyte class as equivalent to our immune class and collapsed the remaining classes (neutrophil, epithelial, plasma, eosinophil, connective, mitosis) into non-immune. Since HoVer-NeXt is also the model we use for nucleus identification, any comparisons are based primarily on the immune classification ability.

**SEQUOIA**: a published set of cancer-type specific models that predict bulk gene expression directly from H&E whole-slide images (25) using foundation model features. We applied it to the relevant cohorts using UNI (57) patch features reduced to a slide-level k-means cluster representation, with the publicly released TCGA-KIRC checkpoints; per-slide expression was averaged across the five cross-validation fold models (gevaertlab/sequoia-kirc-0 through -4).

From the predicted expression we derived a T-effector score as the mean of the corresponding genes (as described above), and compared it, alongside our H&E immune density, against the measured RNA T-effector. Unlike our cell-counting model, SEQUOIA infers an expression signature directly, so the comparison contrasts the two paradigms for recovering T-effector signal from H&E.

### Whole Slide Image Quantification

Each whole slide image was processed through the following pipeline to compute a per-slide immune score, a measure of immune-cell density in and around the tumor, which were then aggregated to a patient-level score.

**1. Tumor Region Identification:** Tumor regions in WSI were identified using an in-house classification model based on the ViT-B/16 architecture, trained on 124 manually annotated H&E WSI from UTSW. Each 224 × 224-pixel patch (∼112 µm at 20X) was assigned to one of eight tissue classes: background, blood, fat, normal kidney, stroma, tumor, necrosis, or inflammatory tissue. The model was applied across each WSI on a grid with a stride of 64 pixels, with each grid point classified from its surrounding patch, producing a low-resolution regional tissue mask. Tumor region assignments were examined where appropriate by an expert pathologist (PK) and manually corrected as needed. In the sarcomatoid cohort, where the goal was to identify the sarcomatoid architecture of tumor tissue, these regions were likewise manually defined when other tissue architectures were present on the slide.
**2. Tissue and Artifact Segmentation:** Tissue regions and slide artifacts were segmented with GrandQC (58), run as its tissue-detection model followed by the artifact-segmentation model at 5x, with otherwise default settings. GrandQC labels each pixel as tissue, background, or one of several artifact types (tissue folds, foreign objects, pen markings, air bubbles and slide edges, and out-of-focus regions); only the tissue-versus-non-tissue distinction was used.
**3. Peritumor Region Identification:** The region-classifier and GrandQC outputs were combined into a single mask, with each grid cell reset to background when more than 10% of its area was non-tissue by GrandQC and the remaining cells keeping their region-classifier label; on a small number of slides flagged during manual review, where GrandQC tissue and background segmentation was unreliable, the region-classifier tissue calls were retained and only GrandQC artifact regions removed. This yielded three compartments: the tumor region (Step 1), the surrounding non-background tissue, and background. A Euclidean distance transform from the tumor region then assigned each location its distance to the nearest tumor. Non-tumor tissue regions within 1mm of the tumor were then defined as peritumoral.
**4. Nucleus Assignments:** Nuclei in each WSI were detected and classified into cell-type classes by the nucleus-level inference described above. Each nucleus was also assigned to a region by the location of its centroid in the region mask: tumor, peritumor, other tissue, or non-tissue background. Nuclei in non-tissue background were discarded.
**5. Immune Score Calculation:** For each WSI, the immune score was computed as a density, the number of immune nuclei in a region divided by the tissue area of that region (cells/mm²). The primary immune score used the tumor together with its 1 mm peritumoral margin (tumor+peritumor) as the region. For a patient with more than one WSI, the patient-level immune score was the maximum of the per-WSI scores, so that for cohorts with a single slide per patient the patient and slide scores are identical.

### Survival Analysis

Survival endpoints differed by cohort: OS for TCGA-KIRC, and PFS for the UTSW Ipi+Nivo, UTSW Nivo, and IMmotion150 cohorts, with event-free patients censored at last follow-up. For IMmotion150, RECIST-based categories additionally defined responders (complete or partial response) versus non-responders (stable or progressive disease) for the objective response rate and AUC analyses. The score under evaluation, the H&E DL Immune score and, for IMmotion150, also the RNA T-effector score and CD8 IHC, was related to outcome without thresholding using Harrell’s concordance index. For thresholded analyses, patients were stratified by that score two ways, a median split and a quartile split (top quartile versus the remaining three); for each, Kaplan-Meier curves were compared by log-rank test and hazard ratios obtained from univariate Cox proportional-hazards models with 95% confidence intervals. Because IMmotion150 comprised three treatment arms, the median and quartile cutoffs were determined on the full cohort and then applied within the relevant arm. For the IMmotion150 response analysis, objective response rate was compared between immune-high and immune-low groups, and discrimination of the continuous score for response was quantified by the area under the ROC curve. Analyses used the lifelines package; two-sided p < 0.05 was considered significant.

## Data/Code Availability

H&E images for TCGA KIRC can be downloaded from the TCGA GDC portal, while the corresponding gene expression data is available from cBioPortal. The data for the TMA cohort can be downloaded from https://doi.org/10.25452/figshare.plus.19324118. IMmotion150 data, including H&E images, Response, T-effector scores and CD8 IHC levels is proprietary to Roche. The anonymized genomic data from 163 patients who granted informed consent to share such data, are made available by Roche at the European Genome-Phenome Archive (EGA) under accession number EGAS00001002928. All other data is available from authors upon reasonable request. The final H&E DL Immune model, and all code used in the manuscript will be released at the <u>Rajaram Lab’s public GitHub page</u> upon publication.

## Acknowledgements

We thank all the patients who provided tissues that enabled this research project. This grant was funded in part by DOD (KC200285) and in part CPRIT (RP220294). J.B., A.C., and P.K. are supported by NIH (Specialized Program in Research Excellence in Kidney Cancer P50 CA196516).

## Acknowledgments

We acknowledge the patients whose samples/data provided the foundation for this study and are grateful to the Kidney Cancer Program and the Clinical Data Warehouse teams for their support and assistance.

## Captions for Supplementary Figures

**Supplementary Figure 1. Orthogonal validation of H&E-derived immune predictions using independent CD3 and CD8 immunohistochemistry datasets. A)** Representative examples comparing H&E images, corresponding CD3 IHC staining, and H&E DL model generated cell-type prediction masks from an independent tissue microarray (TMA) cohort. Predicted immune cell densities closely mirror the distribution of CD3-positive lymphocytes across diverse patterns of immune infiltration. **B)** Correlation between percentage of cells predicted by H&E DL Immune model and CD3-positive cells measured by IHC positivity in an independent TMA cohort. The strong association demonstrates that the model captures generalized T-cell infiltration despite being trained using CD8-based supervision. **C)** Correlation between H&E DL Immune score and CD8-positive cells in the independent IMmotion150 cohort. The model-derived immune score remains significantly associated with CD8 IHC measurements across an external clinical trial dataset, supporting robustness and generalizability.

**Supplementary Figure 2. Workflow for whole-slide immune quantification and illustration of intratumoral immune heterogeneity. A)** Pipeline for generating the H&E deep learning immune score. Whole-slide images undergo tissue-region classification, exclusion of small tumor regions, and quantification of immune cells within tumor and peritumoral compartments. Outputs were manually reviewed and subsequently used for quantitative and spatial analyses. **B)** Representative whole-slide image illustrating substantial intratumoral heterogeneity in immune infiltration. Distinct regions of the same tumor demonstrate marked differences in immune cell density ranging from immune-desert to highly inflamed phenotypes, and increased peritumor inflammation (arrow in purple area) emphasizing the value of including peritumor or multiple slide evaluation.

**Supplementary Figure 3. Relationship between H&E-derived immune infiltration and transcriptomic immune signatures.** A) Spatial distribution of immune cells density in peritumor region surrounding the tumor boundary. Immune cell density decreases progressively with increasing distance from the tumor–stroma interface beyond 1 mm. B) Correlation between H&E immune score in tumor-restricted area and RNA-derived T-effector score in the UTSW-RNA and IMmotion150 cohorts. C) Heatmap showing Spearman correlations between the H&E immune score and previously published RNA-based immune signatures. Strong positive correlations are observed with T-cell rich and immune activation program signatures, whereas weak or negative associations are observed as expected for non-immune signatures, supporting the biological specificity of the model.

**Supplementary Figure 4. Association of H&E-derived immune score with clinical and pathologic variables.** The H&E DL Immune score model was applied to the full evaluable TCGA-KIRC cohort (n=439 WSIs, one per patient) and compared against prognostic variables: **A)** WHO/ISUP nuclear grade, **B)** TNM stage, **C)** tumor size, and **D)** overall survival. Each panel uses the subset of the 439 with the relevant annotation available (n shown per panel). For survival, patients were stratified by H&E DL Immune score quartiles (c-index=0.48); no significant association was observed. **E)** Comparison of H&E DL Immune scores between primary tumors and metastatic lesions across multiple cohorts, including IMmotion150, UTSW ipilimumab/nivolumab-treated patients, and UTSW nivolumab monotherapy-treated patients. Immune scores were comparable between primary and metastatic sites. **F)**. Concordance of H&E DL Immune scores between matched primary and metastatic tumors from the UTSW cohort. Scores demonstrated strong correlation between paired samples (Spearman ρ = 0.89; concordance correlation coefficient = 0.76), indicating preservation of the immune phenotype during metastatic progression.

**Supplementary Figure 5. Predictive performance of RNA-derived T-effector score, H&E immune score, and CD8 IHC across treatment arms in IMmotion150. A)** Forest plot showing hazard ratios for progression-free survival according to biomarker-defined immune status within individual treatment arms. Hazard ratios and 95% confidence intervals are from univariate Cox models (high vs low, split at the full-cohort median) **B)** Objective response rate (ORR; the proportion of patients with a complete [CR] or partial [PR] response) stratified by high versus low biomarker groups for each marker. CR and PR contributions are shown as stacked bars (Atezo: atezolizumab; bev: bevacizumab).

**Supplementary Figure 6. Model prediction correlates with ICI response in contemporary real-word UTSW Ipi+Nivo cohort**. **A)** Kaplan–Meier analyses comparing progression-free survival among biomarker-defined groups in the UTSW ipilimumab + nivolumab and nivolumab cohorts using median as a cutoff. **B)** Bootstrap distributions evaluating robustness of hazard-ratio and concordance-index estimates.

